# Lineage segregation, pluripotency and X-chromosome inactivation in the pig pre-gastrulation embryo

**DOI:** 10.1101/347823

**Authors:** Priscila Ramos-Ibeas, Fei Sang, Qifan Zhu, Walfred W.C. Tang, Sarah Withey, Doris Klisch, Matt Loose, M. Azim Surani, Ramiro Alberio

**Author notes:** Current address: P.R-I.: Animal Reproduction Department, National Institute for Agricultural and Food Research and Technology, Madrid 28040, Spain; S.W.: Stem Cell Engineering Group, Australian Institute for Bioengineering and Nanotechnology, University of Queensland, Building 75, St Lucia, QLD 4072, Australia. These authors contributed equally to this work (Priscila Ramos-Ibeas, Fei Sang).

## Abstract

High-resolution molecular programs delineating the cellular foundations of mammalian embryogenesis have emerged recently. Similar analysis of human embryos is limited to pre-implantation stages, since early post-implantation embryos are inaccessible. Notwithstanding, we previously suggested conserved principles of pig and human early development. For further insight on pluripotent states and lineage delineation, we analysed pig embryos at single cell resolution. Here we show progressive segregation of inner cell mass and trophectoderm in early blastocysts, and then of epiblast and hypoblast in late blastocysts. We detected distinct pluripotent states, first as a short ‘naïve’ state followed by a protracted primed state. Dosage compensation with respect to the X-chromosome in females is attained via X-inactivation in late epiblasts. Detailed human-pig comparison is a basis towards comprehending early human development and a foundation for further studies of human pluripotent stem cell differentiation in pig interspecies chimeras.

---

Pre-gastrulation embryo development shows broad similarities between mammals, although species-specific differences in early lineage segregation, the establishment of pluripotency, and X chromosome inactivation have been reported^1–3^. Mouse embryos, which are widely used as a model for mammalian development, transit rapidly through early development (E3.5-E5.5, i.e. ~2 days), followed by development of the characteristic cup-shaped postimplantation epiblast. In larger mammals, including humans, non-human primates (NHP) and pigs, there is a protracted developmental period (~6 days) that ends with the formation of a flat bilaminar embryonic disc. Since early post-implantation human embryos are largely inaccessible, we are beginning to investigate relatively more accessible pig embryos. Notably both human and pig embryos evidently form a flat embryonic disc before the onset of gastrulation^4^. Thus, the pig embryo can broaden our understanding of the pre-gastrulation development of large mammals with protracted development.

Segregation of trophectoderm (TE) and hypoblast, and the emergence of pluripotency are well established in mice^5,6^, but require detailed studies in other mammals at the resolution of single cells, as recently reported for *Cynomolgus* monkeys^8^. Potential discrepancies in lineage segregation have however emerged in reports between monkey and human, attributed in part to embryo staging differences^7^. Further studies, including those in other large mammalian species, are therefore highly desirable.

In mouse embryos a distinct transcriptional signature of naïve pluripotency in the inner cell mass (ICM) is replaced by a mature epiblast (EPI) identity, marking a transition through different pluripotent states before gastrulation^8^. Whereas naïve pluripotent stem cells (PSCs) resemble ICM epiblast cells and primed PSCs resemble the post-implantation mouse epiblast, establishment of similar cell lines from non-rodent mammalian species, including humans, has been challenging, suggesting possible biological differences^9,10^. Indeed, spatiotemporal differences in the expression of core pluripotency genes *NANOG*, *OCT4* (*POU5F1*) and *SOX2* have been noted, while expression of *Klf2*, *Prdm14*, and *Bmp4* in mouse embryonic naïve cells are not apparently detected in human embryos^9,11^. By contrast, *KLF17* is expressed in the human but not mouse ICM^9,11,12^. Also, while Jak-Stat3 and WNT signalling are detected in the early mouse ICM^13^, many TGFβ signalling components are present in marmoset, human and pig ICM^11,12,14,15^, indicating that the emergence and establishment of pluripotency in mammals is controlled by different signalling pathways and gene networks. Differences in the mechanisms of X-linked gene dosage compensation in female embryos are also evident^3^. The gene dosage compensation with respect to the X chromosomes in female embryos occurs in pre-gastrulation epiblasts in mouse and rabbits^3,8,16^. Notably, human post-implantation and pig pre-gastrulation epiblasts have not been studied^12,16^.

Here we report lineage segregation, the establishment of pluripotency, and X-chromosome inactivation during the entire peri-gastrulation period in the pig embryo using single cell RNA-seq (scRNA-seq). This comprehensive analysis provides new understanding of the developmental trajectories of early embryonic cells in the pig, which shares similarities with the early human development, and other mammals with similar construction.

## Results

### Progressive lineage segregation in pig embryos

First, we set out to generate a single-cell transcriptome profile of early *in vivo* pig embryo development, from four pre-implantation stages: morula (M; embryonic day (E) ~4-5), early blastocyst (EB, ~E5-6), late blastocyst (LB, ~E7-8), and spherical embryo (Sph, ~E10-11)^17^ (Figure 1a), and obtained 220 single-cell transcriptomes from 28 embryos (Supplementary Fig. 1, Supplementary Table 1). Unsupervised hierarchical clustering (UHC) (15,086 genes) grouped the cells according to their developmental stage and specific lineages based on known markers (Figure 1b).

**Figure 1.**
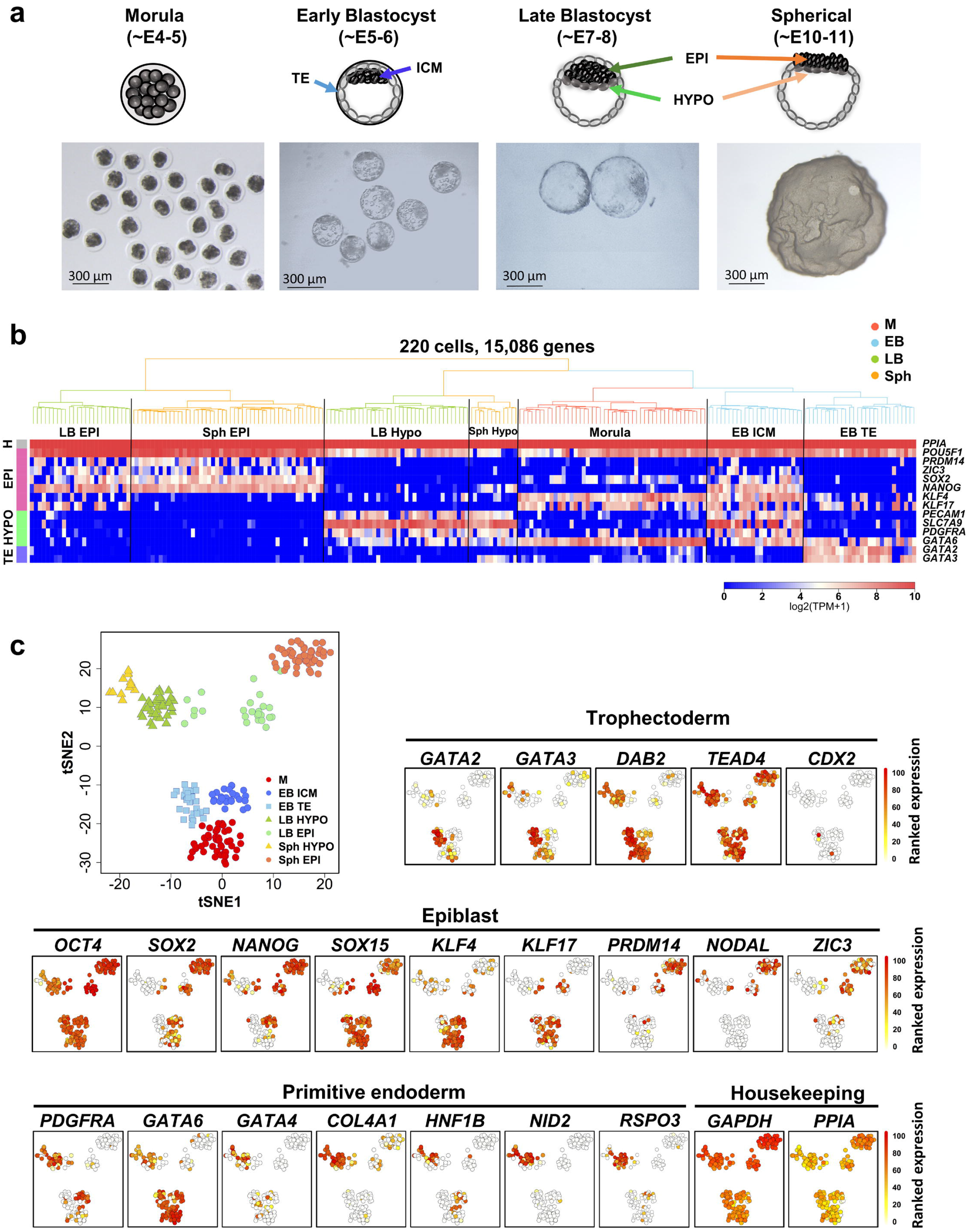
Lineage segregation in pig pre-implantation embryos. **a,** Pig pre-implantation embryos collected for scRNA-Seq **b,** Unsupervised hierarchical clustering (UHC) with all expressed genes (15,086 genes), with a heat map of expression levels of lineage-specific markers. Colors in dendrogram indicate developmental stage. **c,** t-SNE plot of all cells, indicated by colors and shapes for different embryonic days and lineages. Lineage specific genes are shown in t-SNE plots; a gradient from white to red indicates low to high expression. M: morula, EB: early blastocyst, LB: late blastocyst, Sph: spherical embryo, Epi: epiblast, Hypo: hypoblast, ICM: inner cell mass, TE: trophectoderm.

Dimensionality reduction provided a clear visualization of lineage segregation during development (Figure 1c). The morula group showed expression of *OCT4* (*POU5F1*), *SOX2*, and *KLF4*, but not *NANOG*, while early blastocyst (EB) cells segregated into two lineages: ICM cells expressing *NANOG* and *SOX2*, and TE cells with *GATA2*, *GATA3*, and *DAB2*. Expression of *CDX2* was seen in a few TE cells at this early stage^15,18^, but *OCT4* expression was seen in all cells, consistent with observations in human and monkey blastocysts^2,19^. There was evident expression of pluripotency genes; *SOX15*, *KLF4* and *KLF17* in the ICM and EPI, as in human epiblast cells. Expression of some of these genes was also seen in pig TE and hypoblast (HYPO).

We identified 708 differentially expressed genes (DEG) between ICM and TE (Supplementary Figure 2a and b, Supplementary Table 2). While *GATA2* and *GATA3* were the two top-ranked genes in TE of early blastocysts, other TE markers reported in mouse and human such as *ANXA6* and *TEAD1* were identified for the first time in the pig. Notably, we found upregulation of both HYPO and EPI markers in the ICM (Supplementary Figure 2b, Supplementary Table 2). Further interrogation of ICM cells by Principal Component Analysis (PCA) of all genes and highly variable genes (Supplementary Figure 2c and d, respectively) did not separate the cells into discrete populations. Analysis of highly variable genes (HVGs) in a subset of cells separated along PC1 did not show a distinct EPI or HYPO expression signature based on high-confidence markers^7^ (Supplementary Figure 2e). Mutually exclusive segregation of EPI and HYPO became evident first in cells of LB and Sph embryos (TE was excluded from these stages) (Supplementary Figure 2 c and d). Expression of *SOX2*, *NANOG*, *PRDM14*, and *NODAL* was observed in EPI, whereas expression of *PDGFRa*, *GATA4*, *GATA6*, *COL4A1*, *NID2* and *HNF1B* was detected in HYPO (Figure 1c). Comparison between EPI and HYPO in LB and Sph identified 1810 and 1916 DEGs, respectively. Known EPI genes up-regulated in both stages included *SOX15*, *ZIC3*, *FGF19*, *SALL2*, and the HYPO genes *PITX2*, *PECAM1*, *DAB2*, *FN1* (Supplementary Figure 2a and b, Supplementary Table 2). These results show that TE and ICM in the pig embryo segregate in the early blastocyst, whereas at this stage, HYPO and EPI genes are coexpressed in the ICM; these cells resolve into discrete cell lineages in late blastocysts.

### Signalling pathways controlling lineage segregation in the pig embryo

Gene Ontology (GO) enrichment and Kyoto Encyclopaedia of Genes and Genomes (KEGG) pathway analyses indicated that PI3K-Akt and Jak-STAT signalling pathways were overrepresented in ICM and TE of early blastocysts (EB), but the WNT signalling pathway was enriched only in ICM cells. In later stages, PI3K-Akt was over-represented in HYPO and MAPK signalling in EPI. Components of the TGFβ pathways were expressed in both EPI and HYPO (Supplementary Figure 2b, Supplementary Table 2).

For elucidating functional roles of these signalling pathways during lineage segregation, we cultured *ex vivo* pig embryos in the presence of selective inhibitors and determined the impact on lineage allocation by immunofluorescence (IF). In controls, NANOG was absent in morulae, but detectable in most ICM cells from EB (n=9), which were also positive for SOX2 (Figure 2a). Expression of SOX17 was first observed in a subset of NANOG positive (NANOG+) cells (47.14%, n=4) in the ICM of EB, which became gradually restricted to a small group of cells in the ICM of mid-blastocysts (MB, ~E6-7, 16.78%, n=6). By the late blastocyst (LB) stage (~E7-8), two mutually exclusive groups of NANOG+ and SOX17+ cells (n=5) were identified in the ICM (Figure 2a and b).

**Figure 2.**
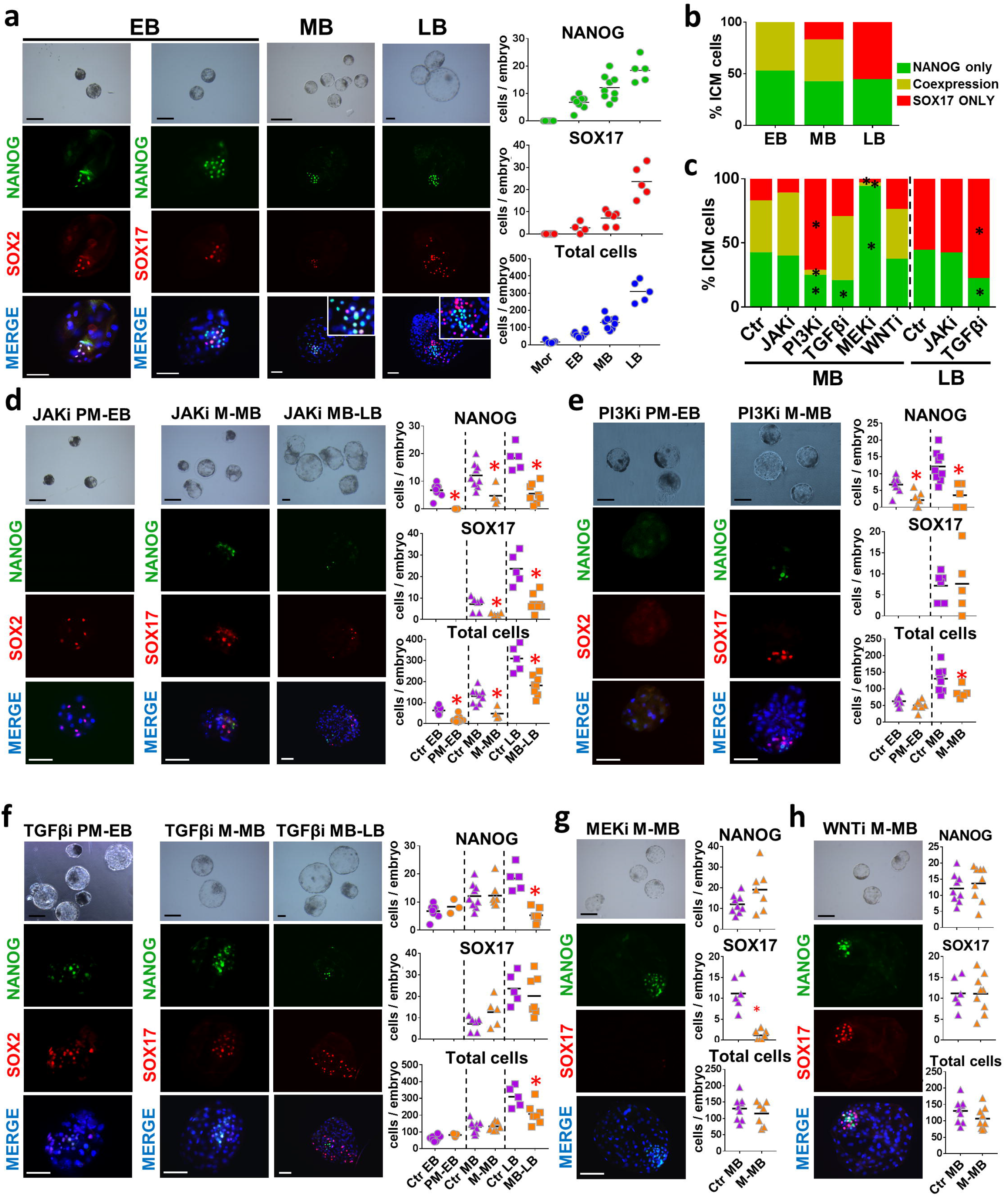
Signaling pathways involved in segregation of lineages. Bright field and IF staining for indicated markers; embryos were counterstained with DAPI (merge). Scatter dot plots of NANOG-, SOX17-positive cells and total cell numbers (black bar indicates mean) of control pig early blastocysts (EB), mid-blastocysts (MB) and late blastocysts (LB). **b**, Bar charts indicating percentage of ICM cells expressing indicated markers in control embryos and **(c)** after different treatments. **d**, Bright field and IF staining of indicated markers in embryos of different stages. Scatter plots show proportion of cells stained for the indicated markers in embryos treated with JAKi: 10 μM AZD1480; **(e)** PI3Ki: 10 μM LY294002; **(f)** TGFpi: 20 μM SB431542; **(g)** MEKi: 10 μM PD0325901; **(h)** WNTi: 3 μM IWP2. PM: premorula, M: Morula, EB: early blastocyst, MB: mid blastocyst, LB: late blastocyst. For **d-h**: * p ≤ 0.05, Mann-Whitney test. For **b** and **c**: * p ≤ 0.05, Two-way ANOVA. Scale bars: 50 μm.

Having established the sequence of NANOG and SOX17 expression, we used these markers to investigate specific signalling pathways. We first looked at Jak-STAT signalling, since it was highly represented in all cells of the EB (Suppl. Figure 2b). Pig embryos cultured from pre-morula (PM, 4- to 16-cells) to EB with a JAK1/2 inhibitor (AZD1480, n=8) had no visible ICM, showed no NANOG expression and a significantly reduced cell number (Figure 2d). While a small number of scattered SOX2+ cells were observed, they were however not organized into an ICM, unlike in control embryos (Figure 2a). In MB (n=4) and LB (n=8) the number of NANOG+, SOX17+ and total cells were reduced, but the relative proportion of NANOG and SOX17 cells in the ICM was unaffected (Figure 2c and d). Thus, pharmacological inhibition of Jak-STAT inhibition affects all lineages at all stages and prevents NANOG expression in the early ICM, but does not influence EPI/HYPO segregation.

We also looked at the role of the PI3K-Akt signalling pathway previously identified in mouse pre-implantation embryos^20^, since we found it enriched in pig EB KEGG terms. Embryos treated with the PI3K inhibitor LY294002 from PM to EB (n=7) and from M to MB (n=5) developed small blastocysts with reduced numbers of NANOG+ cells compared to controls (Figure 2c and e). The total cell number was also reduced, suggesting a role of this pathway in TE development, consistent with our scRNA-Seq analysis (Supplementary Figure 2b). Next, we investigated the TGFβ pathway, which was previously reported during EPI development in human and pig ^11,21,22^. Inhibition of TGFβ signalling in human embryos affects the number of NANOG+ and SOX17+ cells, but there is no effect on lineage segregation^6^. The presence of SB431542 (20 or 40 μM) from PM to EB (n=3) and M to MB (n=7) did not affect embryo development. In contrast, embryos treated from MB to LB (n=7) showed a significant reduction in NANOG+ cells, but the number of cells expressing SOX17 was unaffected (Figure 2c and f). These results indicate that TGFβ is not required for the activation of NANOG, but is necessary for maintaining its expression in the pig epiblast. In human, pig and cattle, inhibition of FGF signalling with a MAPK/ERK kinase inhibitor (PD0325901; 0.4-1 μM) does not abolish the expression of hypoblast markers^23–25^, in contrast to mouse and rabbit embryos where it prevents hypoblast formation^6,26^. As our scRNA-Seq data shows expression of MAPK pathway genes in LB EPI cells, we tested the effect of the MEK inhibitor PD0325901 at high concentration (10 μM), based on previous results with cattle blastocysts^27^. MEK inhibition from M to MB (n=7) significantly reduced the number of HYPO cells resulting in <3 SOX17+ cells/embryo, with an apparent shift towards NANOG+ cells in the ICM (Figure 2c and g). This indicates that MAPK inhibition restricts the expansion of the pig HYPO, but it does not prevent the activation of SOX17 in some cells. Lastly, since regulation of the canonical WNT pathway was a significantly up-regulated GO term in the ICM, we cultured pig embryos with the tested WNT inhibitor IWP2. No reduction in hypoblast segregation nor the total cell number was observed following WNT inhibition from M to MB (n=9) (Figure 2g, h); similar observations were reported for mouse embryos^14^.

### Emerging naïve pluripotent cells and their transition to a primed pluripotent state during epiblast maturation

We next sought to determine how the emergent pluripotent cells (ICM) of EB compare to early (LB) and late EPI (Sph). In a three-dimensional PCA plot, cells grouped as two main clusters: M/ICM and LB/Sph EPI cells (Figure 3a). We detected a biphasic profile of pluripotent gene expression, with high expression of naïve pluripotency genes in M/ICM, and gradual down-regulation of these markers in EPI cells (LB/Sph). Concomitantly with the decrease in naïve markers, there was an up-regulation of primed pluripotency genes in LB/Sph EPI (Figure 3b). Essential differences in gene expression were noted in the pig compared to observations in the mouse^28^; while *OCT4* and *SOX2* expression were maintained along all pluripotent stages, expression of *NANOG* was first observed in the ICM and remained high in LB and Sph EPI. The naïve pluripotency markers *KLF4*, *KLF5*, *KLF17*, *TFCP2L1*, *ESRRB* and *TBX3*, were detected in M and ICM and decreased, or even ceased in LB and Sph EPI. The exception was *PRDM14*, which followed the opposite trend. By contrast, primed pluripotency markers *NODAL*, *DNMT3B*, *SALL2* and *SFRP2* were up-regulated in LB and Sph EPI. Continued expression of pluripotency markers and absence of lineage commitment gene expression (*MIXL1*, *FOXA2*, and *T*) in Sph EPI indicated a protracted exit from pluripotency (about six days) in the pig.

**Figure 3.**
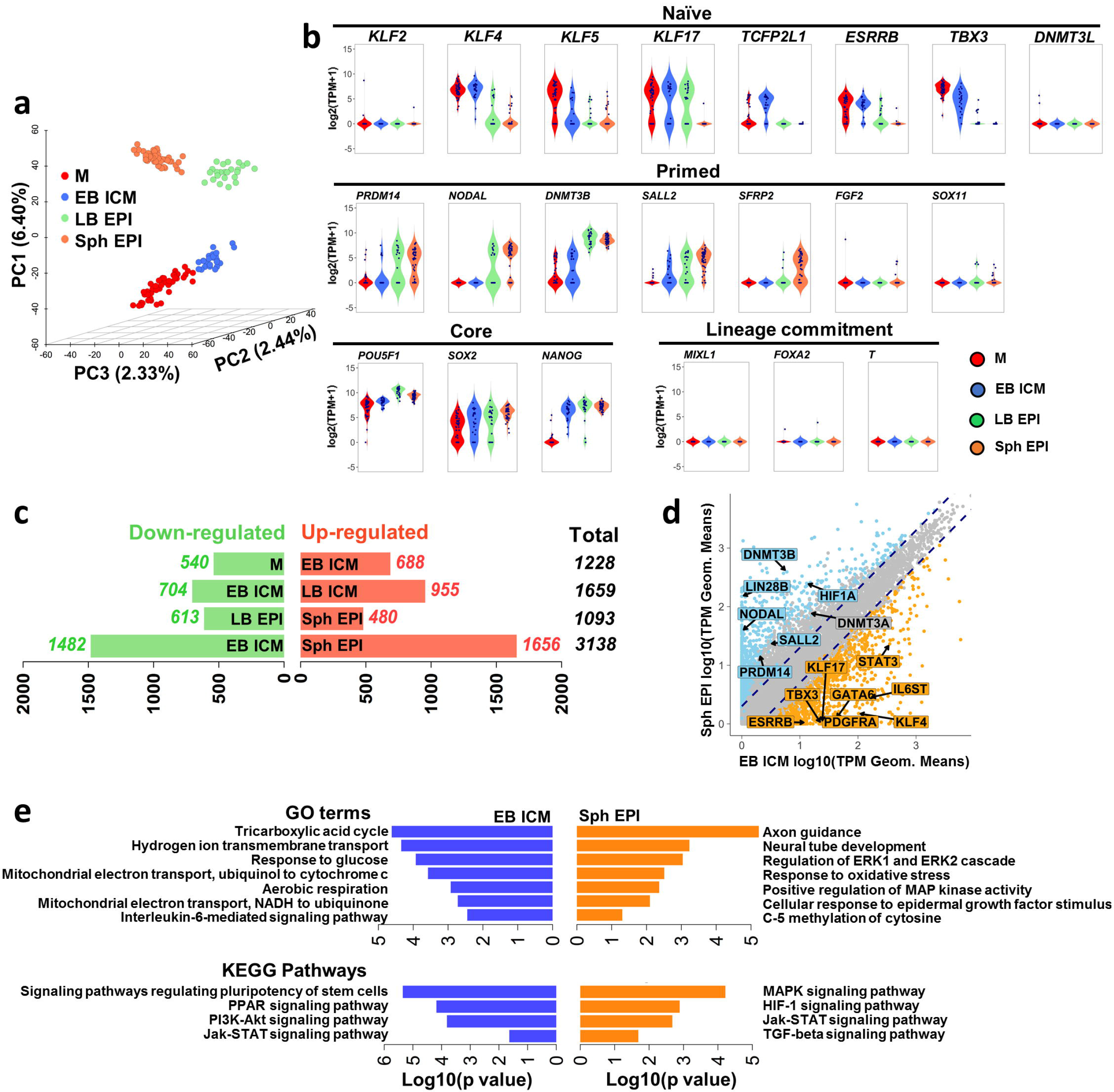
Transitions of pluripotent states in the pig. **a,** Principal component analysis (PCA) of the pluripotent lineages. **b,** Violin plots of the expression of selected genes during the transition of pluripotent lineages. **c,** DEGs during the transition of pluripotent lineages. Red and green bars indicate up- and down-regulated genes, respectively by pair-wise comparisons as indicated. **d,** Scatter-plot of the average gene expression levels between EB ICM vs. Sph EPI (◻1 fold change flanking diagonal lines). Orange: up-regulated, blue: down-regulated; log_10_ (TPM geometric means), key genes are annotated. **e,** Significant Gene Ontology terms and KEGG pathways in DEGs in the pair-wise comparison are indicated.

We used K-means clustering to group genes with similar expression profiles (Supplementary Figure 3a). Genes highly expressed in morulae and ICM cells (cluster 5, 15, 20, 24, 25) include naïve pluripotency markers, members of the Jak-STAT pathway, *TET2*, and components of the Polycomb Repressive Complex 2 (PRC2) *EZH2* and *EED*. Genes up-regulated in LB and Sph EPI (cluster 6, 11, 16, 22) include primed pluripotency markers, DNA methyltransferases, genes indicative of glycolytic metabolism and TGFβ signalling. The transition from naïve to primed pluripotency was further evidenced by the 3138 DEGs between ICM and Sph EPI (Figure 3c, d, Supplementary Table 3). GO enrichment and KEGG pathways analyses between these stages showed that PI3K-Akt, Jak-STAT and Interleukin-6-mediated signalling pathways were upregulated in ICM cells (Figure 3e, Supplementary Figure 3b).

Mouse ICM cells express LIF receptor (LIFR)^29^ and glycoprotein 130 (also known as IL6st)^30^ which bind LIF secreted by the neighbouring TE^31^. However, *LIF* expression was not detected in the pig dataset, consistent with a previous report^32^. Instead, *IL-6* was detected in M and EB TE cells. Similarly, *IL6ST* and IL-6 receptor (*IL6R*) expression were mainly detected in ICM cells (Supplementary Figure 3c). These pathways are down-regulated in LB EPI, and instead, MAPK and TGFβ signalling pathways components become highly expressed. Interestingly, no significant changes in signalling pathways affecting pluripotency were observed between early (LB) and late (Sph) EPI, indicating that the primed EPI stably maintains its properties over ~6 days (Figure 3c, Supplementary Figure 3b).

### Expression of specific surface markers in different pluripotent stages

We sought to identify novel pluripotency markers in the pig, by comparing our dataset with the cell surface protein atlas^33^. naïve and primed pluripotency surface markers in human^34^, such as CD130 (IL6ST) and CD24, were not lineage-specific in the pig embryo (Suppl. Fig. 4a). Instead, we found CD247 primarily marking the ICM and LB EPI, while CD90 (THY1) was present mainly in LB and Sph EPI (Supplementary Figure 4a). Notably, CD200, CD79B and CD83 were specifically expressed in late epiblast cells and could constitute primed pluripotency cell surface markers in the pig. Candidates for new naïve markers were CD200R1, expressed only in M and ICM cells, and CD244, expressed exclusively in ICM. We confirmed the expression of CD244 by IF, which unexpectedly showed nuclear localization within a subpopulation of SOX2+ cells in the ICM (n=12) of EB. Some of these cells also showed SOX17 co-expression. By the MB stage, CD244 was almost undetectable (n=5), consistent with scRNA-seq data showing down-regulation of this marker in late blastocysts (Supplementary Figure 4b).

### Distinct metabolic and epigenetic programs govern the transition of pluripotent states

A shift towards glycolytic metabolism and reduced mitochondrial activity is associated with the development from naïve to primed pluripotency in mouse and human PSC^35,36^; this metabolic switch has also been described in the mouse epiblast^35^. DEG and GO term analysis between ICM and Sph EPI cells suggested a metabolic switch during the transition of pluripotent states in the pig embryo (Figure 3d, e). Notably, *ESRRB* and *STAT3*, which stimulate oxidative phosphorylation (OXPHOS) during maintenance of naïve pluripotency^37,38^, were up-regulated in M and ICM, but were later down-regulated in LB and Sph EPI. Enzymes involved in the tricarboxylic acid (TCA) cycle and OXPHOS, such as *IDH1*, *ACO2* and *UQCRC2* followed the same trend, as well as *EGLN1*, which prevents HIF1a stabilization and is down-regulated in primed pluripotent cells^39^ (Figure 4a; Suppl. Fig. 5). *LIN28A* and *LIN28B* maintain low mitochondrial function in primed pluripotent cells^36,40^, and *MYC* binds to the *LIN28B* locus and potentiates glycolysis^41^. These genes were up-regulated in pig EPI. A similar expression pattern was noted for *HIF1α*, a hypoxia-inducible factor up-regulated during the transition from naïve to primed state^35^, concomitantly with the up-regulation of downstream enzymes *HK1*, *GBE1*, *PGM1*, and *PYGL*, required to convert glucose to glycogen. Finally, the glycolytic enzymes *LDHA^35,42^* and metabolite trasnporter *UCP2*, which limit pyruvate oxidation and facilitate glycolysis^43^, were also up-regulated in EPI cells (Figure 4a; Suppl. Fig. 5). We also detected a reduction in expression of electron transfer complex IV (cytochrome c oxidase) genes (11/20 genes) during the maturation of the epiblast (Figure 4b), suggesting a reduction in mitochondrial metabolism^35^.

**Figure 4.**
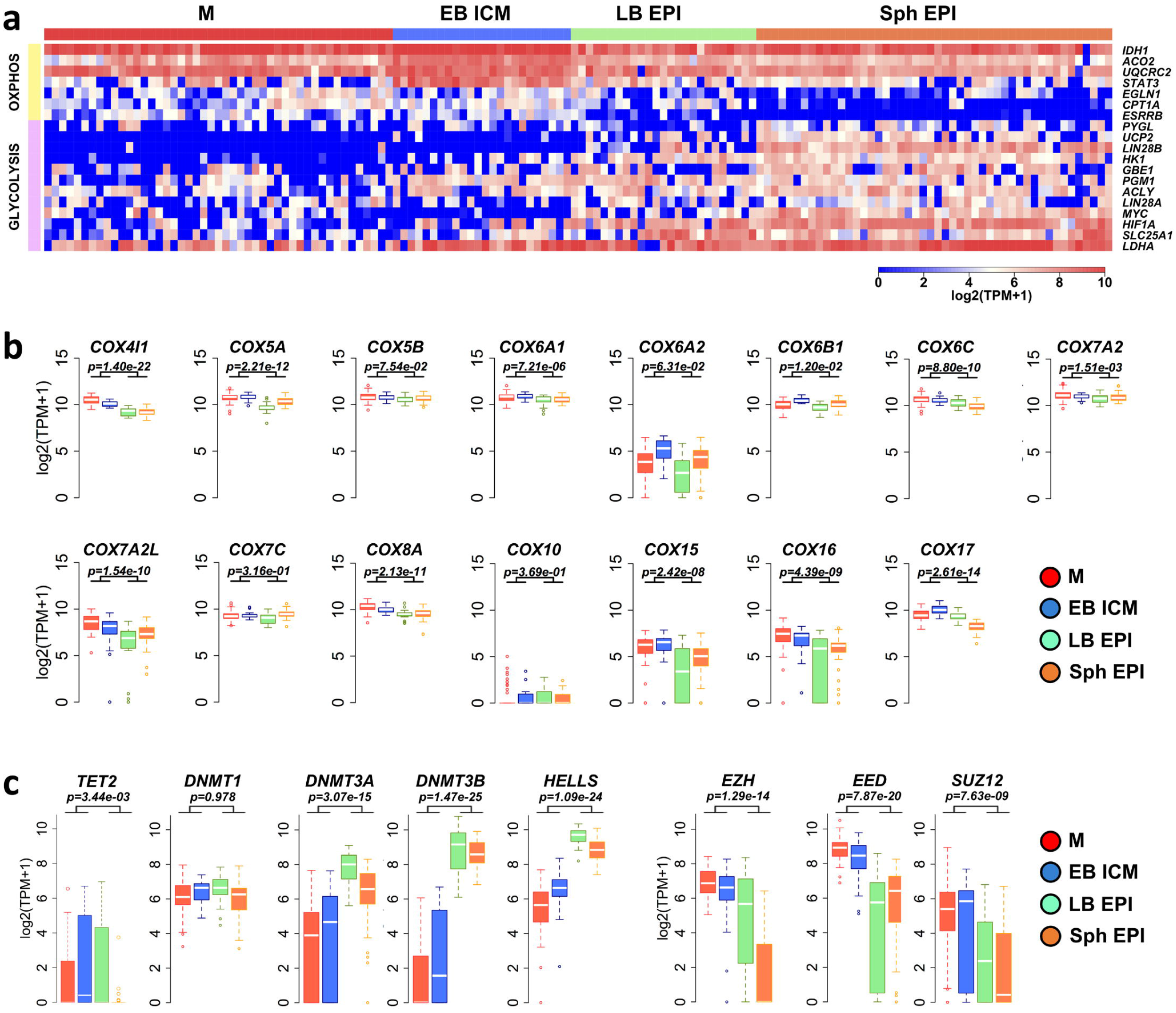
Metabolic and epigenetic transition during changes in pluripotent states. **a,** Heatmap of selected genes involved in OXPHOS and anaerobic glycolysis in pluripotent lineages. **b,** Box plot showing expression of electron transport complex genes and **(c),** genes involved in epigenetic modifications. Two-sided Wilcox test. M: morula, EB: early blastocyst, LB: late blastocyst, Sph: spherical embryo.

We observed down-regulation of the fatty acid transporter to the mitochondria *CPT1A* and a concomitant increase of critical fatty acid synthesis genes *SLC25A1* and *ACLY* in EPI cells compared to ICM; this is in agreement with previous reports indicating accumulation of long-carbon-chain lipids during the conversion from naïve to primed pluripotency in mouse and human^39^ (Figure 4a).

Epigenetic modifications are highly responsive to metabolites derived from pathways such as the TCA cycle or glycolysis, in particular, DNA methyltransferases (DNMT), histone acetyltransferases and histone methyltransferases ^44^. GO terms related to *de novo* DNA methylation were up-regulated in EPI cells (Figure 3e, Suppl. Fig. 3b). Accordingly, the expression of *DNMT3A*, *DNMT3B* and *HELLS*, required for *de novo* DNA methylation^45^, significantly increased in LB and Sph EPI. Concomitantly, *TET2* was down-regulated in the late EPI (Figure 4c).

The core components of PRC2 complex *EZH2*, *EED* and *SUZ12* repress developmental regulators through establishing trimethylation of lysine 27 in histone 3 (H3K27me3) modification^46^, preventing differentiation of PSCs^47^. These genes were expressed at all stages harbouring pluripotent cells in the pig embryo, while expression of *EZH2* and *EED* was down-regulated in primed pluripotent stages (Figure 4c), similar to previous observations in pig epiblasts^48^ and human PSCs^39^. Hence, two populations of pluripotent cells with distinct metabolic and epigenetic profiles exist in the early pig embryo.

### Dosage compensation of X chromosome during pig embryo development

To establish the gender of each cell/embryo, the cumulative level of Y chromosome gene expression was established (Supplementary Figure 6a-c). The female-to-male ratio of X-chromosome (XC) gene expression was higher in females from morula to LB in all embryonic lineages, suggesting lack of dosage compensation. However, in Sph EPI, XC gene expression was comparable to that of autosomes in all embryos, indicating the occurrence of dosage compensation (Figure 5a). Analysis of XC gene expression relative to autosomes at the single-cell level showed uniformity between male and female cells and confirmed dosage compensation in Sph EPI (Figure 5b). A chromosome-wide analysis of female-to-male ratio showed a progressive reduction in gene expression along the whole chromosome with some areas maintaining high ratios of expression at the spherical stage (Figure 5c). In agreement with the dynamics of dosage compensation, *XIST* expression was detected in most (81.8%) female cells in the EPI of Sph embryos, with only sporadic expression of *XIST* in some male cells (Figure 5d).

**Figure 5.**
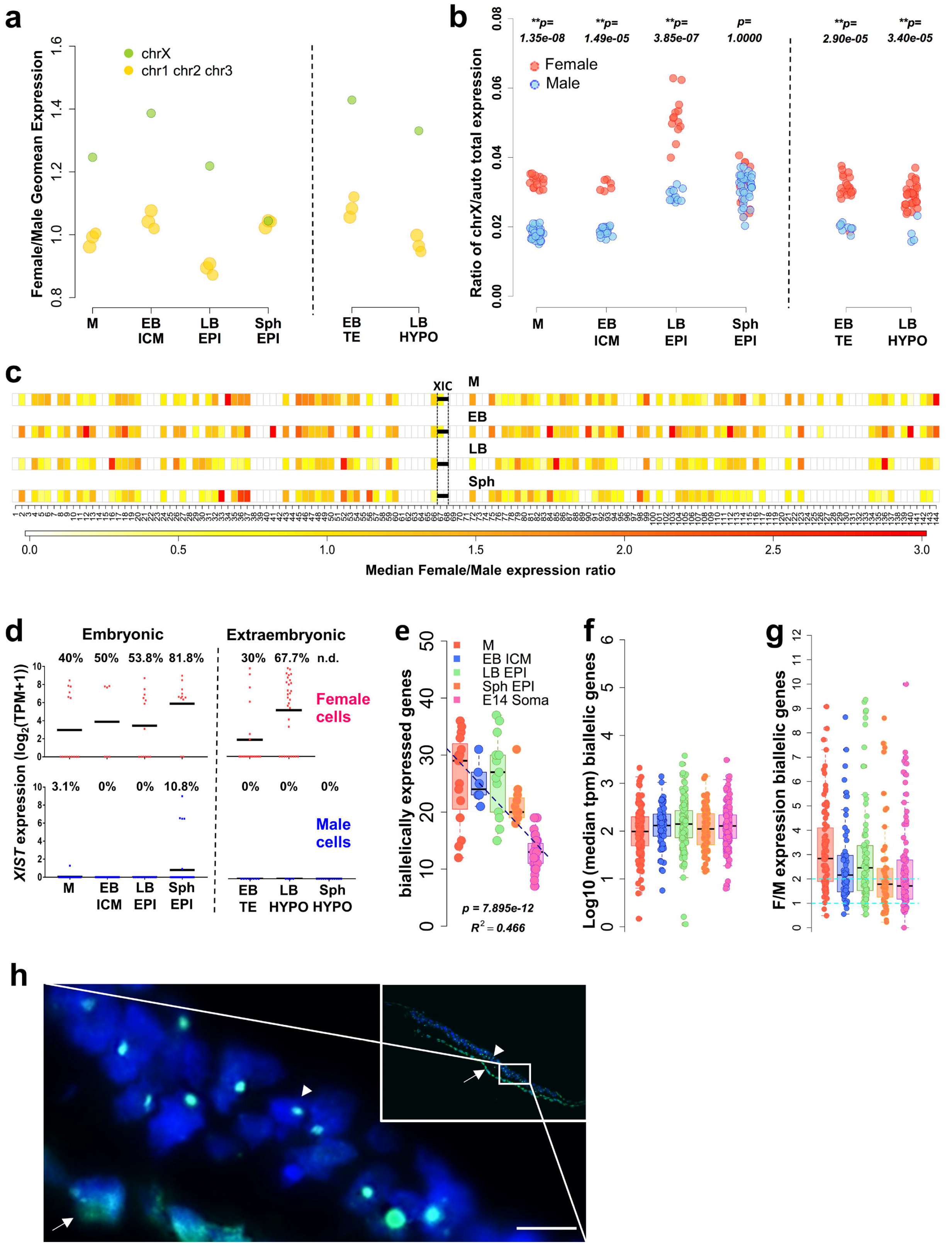
Dosage compensation for the female X chromosome. **a,** Ratio of gene expression between female and male embryos for the X chromosome vs. autosomes 1, 2 and 3. **b,** Proportion of total expression levels of the X chromosome relative to autosomes at the single cell level. **c,** Female to male expression average along the X chromosome. XIC: X-inactivation center. **d,** *XIST* expression level in male and female cells. Percentage of cells with TPM > 1 is shown. **e,** Number of biallelically expressed genes in each cell at different stages of development. **f,** Median expression of biallelic genes. **g,** Female to male ratio of expression of genes biallelically expressed in females. **h,** IF staining of H3K27me3 merged with DAPI in sectioned spherical female embryo. Arrow indicates hypoblast and arrowhead marks the epiblast. Inset shows a low magnification image of the embryonic disc. Scale bar: 10 μm. M: morula, EB: early blastocyst, LB: late blastocyst, Sph: spherical embryo.

To investigate the mechanism of dosage compensation, we analysed XC expression at an allelic resolution, quantifying the expression of single nucleotide variants (SNV) within each cell for a reference or an alternative allele. As expected, SNVs were not found in male cells, consistent with the presence of a single XC (Supplementary Figure 6d). Notably, there was a sharp decline in the number of biallelically-expressed genes in spherical EPI. The lowest level was detected in female mesoderm cells from E14 embryos, where we detected an inactive XC (Fig. 5e; Suppl. Fig. 6e), which served as a somatic cell control. This result indicates that dosage compensation at the spherical stage is attained by inactivation of one XC. To gain a better understanding of the X-inactivation process, we analysed the median expression of biallelically-expressed genes. No median reduction in biallelic gene expression was detected *en route* to dosage compensation (Sph EPI) (Figure 5f). The female/male ratio of biallelically-expressed genes was close to 2 in the stages, which showed dosage compensation (Figure 5g). This result suggests that “dampening” of X-linked gene expression does not precede dosage compensation. To confirm the inactivation of one X chromosome in the epiblast of female spherical embryos, we analysed Histone H3 lysine 27 trimethylation (H3K27me3), which accumulates in the inactive X^49,50^. A clear single focal enrichment of H3K27me3 was detected in the nuclei of epiblast cells in female spherical embryos (Figure 5h), similar to what is observed in mesodermal cells (Supplementary Figure 6e). In contrast, no H3K27me3 foci were found in female LB cells, consistent with the lack of XCI (Supplementary Figure 6e).

## Discussion

We revealed the molecular features of early lineage segregation, pluripotency and X inactivation during development of early pig embryos. Our study provides the basis for comparisons with human and mouse development, and for insights in conservation and divergence of early mammalian development.

Segregation of the first three lineages occurs progressively during preimplantation development, starting with the TE and the ICM in early blastocysts. High levels of *GATA2* and *GATA3* expression detected in early pig TE cells conform to the observations in early human and *Cynomolgus* monkey TE^2,19^. By contrast, *Cdx2* expression is among the earliest markers in the mouse TE^51,52^. ICM cells of early pig blastocysts co-express EPI (*NANOG*, *SOX2*) and HYPO (*PDGFRa*, *SOX17* and *GATA6*) markers, but during the mid-/late blastocyst stage EPI and HYPO lineages become definitively segregated. Our analysis shows that hypoblast cells segregate from a population of NANOG/SOX17+ cells, indicating that ICM cells are bi-potent, able to give rise to mature EPI and HYPO, as shown in mouse^5^, human^7,23^ and monkey^2^.

Pathway analysis revealed Jak-STAT and PI3K-Akt signalling enrichment in TE and ICM of early embryos, and in HYPO of late blastocysts. Jak-STAT signalling is important for the proliferation of appropriate numbers of cells in each of the compartments of the blastocyst, while PI3K-Akt signalling affects total cell numbers and EPI expansion, without affecting segregation of HYPO. The Jak-STAT pathway is an effector of multiple ligand/receptor interactions including members of the IL-6 family, such as LIF, GCSF, and IL-6. Previous studies showed LIFR expression in the TE of late pig blastocysts^53^, which is essential for the development of this lineage^24^. Although there is expression of LIFR in some ICM and TE of early blastocysts, there is no expression of LIF; either in any of the cells of the blastocyst, or in the maternal endometrial cells at this stage^54^, suggesting that LIF signalling does not have a significant role in early pig embryos. Instead, expression of IL-6 in morulae and TE cells, and IL-6R and the co-receptor IL6ST (also known as GP130) in ICM cells, suggests that IL-6 likely activates the Jak-STAT pathway by binding to its cognate receptor. Consistent with this, supplementation of IL-6 during derivation of pig ESC promotes proliferation of blastocyst outgrowths^32^, and could potentially support the derivation of naïve pig ESCs. Notably, IL-6/IL6ST can support derivation of germline competent mouse ESC^55,56^, indicating that this pathway may be conserved in mammals, whereas the role of the LIF/LIFR in pluripotency may be evolutionarily divergent in mice and rats.

Signalling via MEK is important for hypoblast formation in mouse^6^ and rabbit^26^; though this does not seem to be the case in human^23^, marmoset^14^, pig^24^, and cattle^25^. Only when using high concentrations of MEK inhibition we detected a drastic decrease in SOX17 expression, as reported in cattle^27^, suggesting that alternative pathways may be supporting HYPO segregation in large mammals.

Our study reveals that TGFβ signalling is critical during the expansion of the epiblast between MB-LB transition, but not in EPI/HYPO segregation, consistent with previous reports ^24,25^. TGFβ signalling is needed for hESC self-renewal^57^, and inhibition of this pathway affects NANOG expression in human and marmoset blastocysts^11,14^. Similarly, NANOG expression in pig embryos is also affected by inhibition with SB431542. Furthermore, we show high expression of TGFβ components in EPI cells compared to ICM, suggesting that this pathway becomes active in advanced embryos, pointing to a critical role of TGFβ during the expansion of the epiblast.

Analysis of pluripotent embryonic cells revealed a state consistent with naïve pluripotency in morula and ICM cells (made of ~ 10-15 cells) of EB (E~5-6), and a state of primed pluripotency in the EPI of LB (~E7/8) and Sph (~E10/11) embryos, which coincides with an expansion of the epiblast from ~25 cells in LB to more than ~180 in Sph. The rapid transition (about 1 day) from naïve to primed states suggests that naïve cells are unlikely to self-renew. However, the protracted period of primed pluripotent state in the pig embryo offers opportunities for their isolation and expansion as self-renewing cells *in vitro*. Indeed, pig pluripotent cell lines with primed characteristics have been reported^21,58–60^, but not of those with characteristics of naïve pluripotency; the latter may require different culture conditions capable of stimulating their proliferation. Differences in naïve pluripotency properties between mouse and other mammals may underlie the difficulties in establishing equivalent cells from the latter in vitro. Pig naïve pluripotency markers include *KLF4/5/17*, *TBX3* and *TCFPL21*, and are consistent with those reported in human^11^ and monkeys^2,14^, which differ from mouse naïve pluripotency, which is characterized by the expression of *Klf2*, *Prdm14* and *Bmp4*. These genes participate in regulating gene expression, epigenetic reprogramming, and cellular signalling, respectively, which highlight potential functional differences in naïveté between mice and larger mammals.

The transition from naïve to primed pluripotent states in the pig embryo is accompanied by a metabolic shift from OXPHOS towards glycolysis, consistent with an increased proliferation rate^61^. This metabolic switch likely provides critical metabolites to promote epiblast expansion, as well as epigenetic remodelling through epigenetic modifications of DNA and histone^43^, a crucial step in preparation for the next major developmental event that is the onset of gastrulation.

Diverse mechanisms exist in mammals for dosage compensation with respect to the XC in females^62^. In mice, imprinted XCI results in inactivation of the paternal XC in early cleaving embryos, followed by reactivation in the ICM of blastocyst (excluding the extraembryonic tissues), and then random XCI in the epiblast^63^. In contrast, there is no imprinted XCI in human and rabbit embryos. Indeed, the expression of *XIST* from both X chromosomes in blastocysts suggests alternative mechanisms of dosage compensation^3^. The ‘dampening’ of X-linked genes from both parental chromosomes as a possible mechanism^15^ warrants further studies^64^. Another report indicated incomplete dosage compensation of a subset of X-linked genes in pig blastocysts^16,65^. Notably, our observations however show XCI in the mature EPI, as demonstrated by the reduction in the number of biallelically expressed X-linked genes, coupled with the appearance of the H3K27me3 mark on the inactive XC.

Our study at the resolution of single cells allows comparisons between species to identify developmental equivalence. Comparison of mouse and pig pluripotent matched stages showed broad developmental alignment, although the developmental time in mice is three times shorter compared to pigs (2 days vs. 6 days). Yet the overall principles of the emergence and establishment of pluripotency are conserved between these species (Figure 6a). Developmental progression showed broad equivalence between morula to epiblast transitions in humans and pigs. Importantly, human embryonic stem cells (hESC) with naïve and primed characteristics grouped closely to human late ICM and EPI cells, respectively, and these also aligned with pig EPI cells (Figure 6b). Our observations may be relevant for understanding events during early human development, as well as for attempts to study specification of hESCs in chimeras with pig embryos as hosts, following their introduction into blastocysts. Hitherto, the reported efficiency of these experiments is very low^66^, perhaps because the hESCs were not *in-sync* with the host pig blastocyts. Developmental synchrony between donor and host is important for efficient chimerism ^67^. We propose that the introduction of primed state hESCs into late pig blastocysts may be a more favourable environment for homing of hESC, and their subsequent development in chimeras.

**Figure 6.**
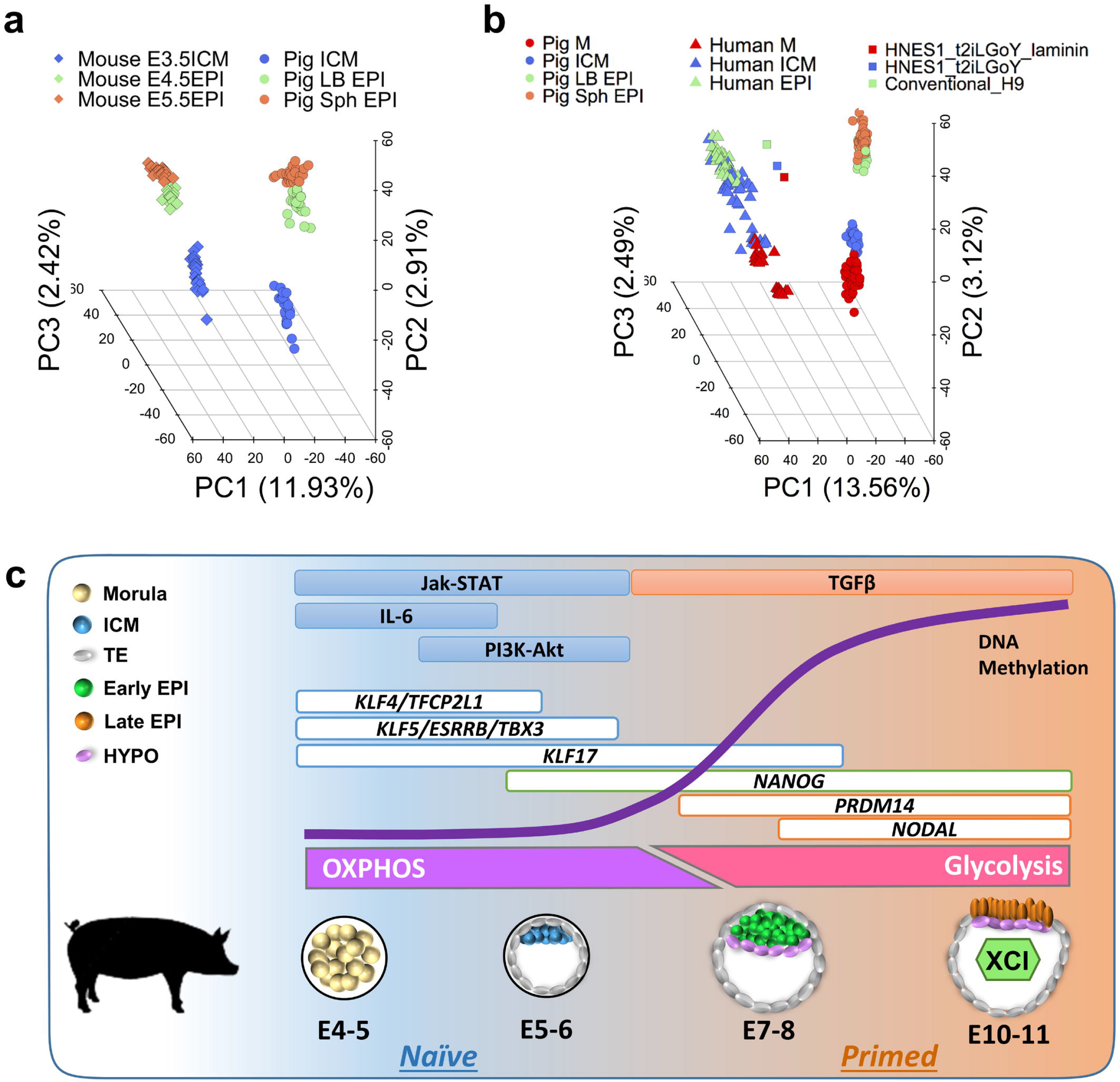
Comparison of pig, mouse and human matched pluripotent states. **a,** PCA of pig and mouse orthologous genes expressed in pluripotent cells. **b,** PCA of pig and human orthologous genes expressed in embryonic cells and hESCs. **c,** Summary of key events in the pluripotent compartment of the pig embryo.

In conclusion, this comprehensive analysis depicts molecular landmarks of pig embryogenesis that provides new insights into embryos with protracted epiblast development (Figure 6c). Furthermore, the shared features of lineage segregation and pluripotency between humans and pigs revealed here will help accelerate research into novel approaches in regenerative medicine, such as the development of interspecies chimeras.

## Methods

### Porcine embryo collection

All of the procedures involving animals have been approved by the School of Biosciences Ethics Review Committee, The University of Nottingham. Embryos at each stage were retrieved from multiple crossbred Large White and Landrace sows (2–3 years old) between days 4 and 11 after artificial insemination. Embryos were flushed from the uterine horns with 30–40 ml warm PBS (supplemented with 1% FCS), washed and transported to the laboratory in N2B27 supplemented with 25 mM HEPES in a portable incubator at 38.5 °C.

### Isolation of single cells for single-cell cDNA preparation

Zona pellucidae were removed using acidic Tyrode’s solution (Sigma) in morulae and early blastocysts, and then embryos were dissociated. Late blastocysts were subjected to immunosurgery to remove the trophectoderm based on previously described procedures^68^. Briefly, embryos were incubated for 30 min in a 1:5 dilution of anti-pig serum (Sigma) in N2B27 medium, washed and incubated for 30 min in 1:5 dilution of complement (Sigma). Embryos were transferred to N2B27 for a few minutes for efficient cell lysis, and then embryonic disks were isolated from the trophectoderm by repeated aspiration with a pulled glass capillary. In spherical embryos, epiblast and hypoblast were manually isolated. Trophectoderm cells were not collected from late blastocysts and spherical embryos.

Single cell dissociation was performed by incubation in TrypLE Express (GIBCO) for 5 minutes at 37 °C and repeated pipetting using very thin pulled capillaries. Individual cells were subsequently transferred to DMEM + 20% FCS to block TrypLE Express and washed in a small drop of PBS-PVP. Single cells were manually collected into PCR tubes to prepare single-cell cDNA libraries following the Smart-seq2 protocol^69^.

Briefly, single cells were lysed by incubation at 72 °C for 3 min in PCR tubes containing four μl of cell lysis buffer, oligo-dT primer and dNTP mix. Reverse transcription and PCR preamplification were carried out with SuperScript II (Invitrogen) and KAPA HiFi HotStart ReadyMix (KAPA Biosystems) respectively according to Picelli *et al.* protocol. PCR products were purified using Ampure XP beads (Beckman Coulter), and library size distribution was checked on Agilent dsDNA High Sensitivity DNA chips on an Agilent 2100 Bioanalyzer (Agilent Technologies). Concentration was quantified using Qubit Quant-iT dsDNA High-Sensitivity Assay Kit (Invitrogen). Samples with more than 0.2 ng/μl, free of short fragments (<500 bp) and with a peak at around 1.5-2 kb were selected for library preparation with Nextera XT DNA Library Preparation Kit (Illumina). Tagmentation reaction and further PCR amplification for 12 cycles were carried out, and PCR products were again purified using Ampure XP beads. Quality of the final cDNA library was analysed on an Agilent high-sensitivity DNA chip. Final cDNA libraries had an average size of 700-800 bp and were quantified using NEBNext Library Quant Kit for Illumina (New England BioLabs) following the manufacturer instructions. Finally, libraries were pooled in groups of 50 with a 2nM final concentration, and DNA sequencing was performed on a HiSeq 2500 Sequencing System (Illumina).

### Data availability and Single Cell RNA-Seq Data

The scRNA-Seq datasets generated during this study are available under GEO accession number: GSE112380. Raw PE reads were trimmed against adaptor sequences by *scythe* (v0.981), and quality-trimmed by *sickle* (v1.33) using default settings. Trimmed reads were directionally aligned to the pig genome (*Sus scrofa* v10.2) by *hisat2* (v2.1.0) with -*know-splicestie-infile* setting to increase mapping accuracy of splicing reads. Uniquely and correctly mapped reads were extracted for the downstream analysis. *htseq-count* was used to count the number of reads aligned to each gene (*Sus scrofa* v10.2 ensembl annotation build 87). Gene expression level was calculated and normalised by Transcripts Per Kilobase Million (TPM).

Low quality cells were filtered out from the dataset to reduce the downstream analysis noise. First, the total number of reads mapped to gene transcripts was calculated for each cell, and those with less than 1 million were removed. Second, the proportion of reads aligned to mitochondrial genes was estimated, as a high proportion suggests poor quality cells^70^. The proportion cut-off was set at 0.5. Only cells of proportions below 0.5 were kept for the next analysis. Third, 4 outlier cells were identified by t-SNE dimensionality reduction. A total of 13,815 out of 22,824 annotated genes were identified in at least 3 cells with TPM > 1.

### Lineage Segregation of Cells

The R package “*scater*” was applied to normalise read counts of genes for each good quality cell with acceptable sequencing coverage. A non-linear approach, t-stochastic neighbour embedding (t-SNE), was used to identify the relations between cells using normalised read counts. Unsupervised hierarchical clustering using all expressed genes as input was conducted on all filtered cells by normalised read counts in log2 scale. The distance method was *euclidean*, and the cluster method was *ward.D2*.

### Lineage Differential Expression Analysis

Pairwise comparisons of single cell differential expressions were performed by SCDE using normalised read counts among four embryo stages. Two-tailed adjusted *p-value* were calculated using cZ scores from Benjamini-Hochberg multiple testing corrections, and followed a normal distribution. Significantly expressed genes were selected with a *p-value* less than 0.05 as the threshold. A heatmap of differentially expressed genes (DEGs) was created with a log2 scale of normalised expression. *Euclidean* distance and default *hclust* were applied to determine the relationships between cells and between genes. Gene Ontology (GO) gene set enrichment analysis with DEGs utilised *goseq* for each pairwise comparison, also with upregulated DEGs and downregulated DEGs separately. GO term annotation was retrieved from the Ensembl database (*Sus scrofa* v10.2 ensembl annotation version 87). Enrichment analysis of biological pathway was performed with DEGs by *gage*. Ensembl gene IDs of DEGs were mapped to NCBI gene IDs for KEGG pathway prior to enrichment analysis.

### Lineage Subpopulation Analysis

Cell lineages were investigated for subpopulation analysis. An outlier EB ICM cell was excluded by PCA based on log2 TPM values of all expressed genes. LB and Sph cells were grouped together for PCA. Two contrasting methods of PCA were applied; one using all expressed genes, and the other based on the highly variable genes (HVGs) only. We used *decomposeVar* to detect HVGs with a loess regression fit model. FDR <= 0.05 was applied as a significance cut-off.

### Inference of Embryonic Sex

Expressions of all the Y chromosomal genes were summed up to determine the sex of each cell. A cell with the total chromosome 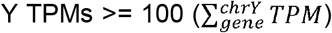 was regarded as a male cell, while 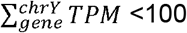 regarded as a female cell.

### Chromosome X dosage compensation analysis

Genes of Chromosome X and three autosomes (chr1, chr2, chr3) were extracted, and the geometric mean TPM of chromosomal expressed genes was calculated for each cell separately. Then the overall geometric mean TPM was obtained for each developmental stage by embryo sex, as well as the total TPM. Each TPM value was incremental by one (TPM+1) for the calculation of geometric mean TPM. Only shared expressed genes between female and male cells were taken into account in the calculation of Female/Male expression ratio for each chromosome. For each cell, the ratio of chrX/auto was inferred by total TPMs, and grouped by embryo sex. Median Female/Male expression ratio was estimated for each stage across the whole chromosome X with 1Mb window.

### Analyses of allelic expression

Trimmed reads were aligned to chromosome X of the pig genome (*Sus scrofa* v10.2) by *hisat2*. Duplicated reads were marked by *picard* (v2.12.1). GATK (v3.8) was used to retrieve allelic read counts for SNVs annotated in dbSNP (build 147). Only validated SNVs (dbSNP flag VLD) were extracted for downstream analysis. SnpEff (v4.3) was applied to annotate called SNVs with *Sus scrofa* v10.2 ensembl annotation (build 87). Low coverage SNVs (< 3 reads) were excluded from the analysis, and we only kept SNVs that occurred at least in two different cells for each stage. The expressions of mono-/bi-allelic genes were estimated based on SNVs in each female cell of each stage.

### Gene clustering by expression profile

Self-Organizing Map (SOM) were used to discover potential structural patterns in highly dimensional and complex datasets by creating 2-dimensional representations. The SOM algorithm was applied to our gene expression profile. The geometric mean TPM of each gene was calculated within each stage. In order to fit the SOM model, the lower TPMs (< 1) were replaced by 1, and the extreme higher TPMs were replaced by 10,000. Similarly, genes of highly similar expression profiles within stages were excluded 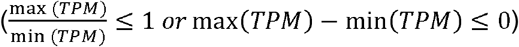. The filtered TPMs were then normalised by SOM(*μ* = 0, *σ* = 1). In total, 25 clusters were created.

### Comparison of pig, mouse and human datasets

In total, 144 pig cells (our study), 83 mouse^8^ cells and 152 human cells retrieved from Petropoulos *et al.*^12^ and re-classified according to Stirparo *et al.*^7^ were included in the comparative analysis. PSC lines cultured under conventional or naïve conditions were used for analysis as described by Stirparo *et al.*^7^. Pig orthologous genes (15,171) were retrieved against human genes from Ensembl database (compara build 87). Expression values were normalised by TPM for PCA analysis. Linear regressions were calculated separately for PC2 and PC3, which contributed to pig and human developmental genes, respectively.

### Embryo treatments with inhibitors and IF

Embryos recovered were incubated in PZM5 culture medium up to morula stage and in N2B27 medium supplemented with 0.3% fatty acid free BSA from compact morula onwards, in a humidified atmosphere at 39 °C and 5% O_2_. The embryos were treated with the following inhibitors and concentrations: 10 μM PD0325901 (Tocris), 20 μM SB431542 (Tocris), 10 μM LY294002 (Selleckchem), 2.5 μM IWP2 (Sigma), 10 μM AZD1480 (Sigma). All treatments were performed for 48h during the indicated time points; from pre-morula (PM) to early blastocyst (EB), from morula (M) to mid blastocyst (MB) and from mid blastocyst to late blastocyst (LB). Inhibitors were dissolved in DMSO and control embryos were treated with DMSO accordingly.

After the treatments, embryos before hatching stage were treated with Tyrode’s acid to remove zona pellucidae. Then, embryos were fixed in 4% paraformaldehyde (PFA) for 15 minutes at room temperature (RT), washed in PBS-1% BSA, permeabilized in 0.2% Triton X-100 for 15 min at RT and blocked in blocking solution (PBS with 0.1% BSA, 0.2% Tween and 10% Donkey serum) for 1 hour at RT. Embryos were incubated overnight at 4°C with the primary antibodies: NANOG (Peprotech, 500-P236, 1:200 dilution in blocking solution), SOX17 (R&D, AF1924, 1:200). After 4 washes in PBS-1% BSA, embryos were incubated in the appropriate secondary antibodies for 45 min at RT, followed by 4 washes in PBS-1% BSA. Finally, embryos were mounted in Vectashield with DAPI.

### Statistical analysis

To evaluate the statistical differences in cell count numbers from individual embryos, probability (p) values were calculated using Two-sided Mann-Whitney test between each treatment and the control. Percentages of contribution of NANOG+ only, SOX17+only and co-expressing cells were evaluated by two-way ANOVA (Dunnett’s multiple comparisons test). Differences were considered significant when p<0.05.

## Acknowledgements

This project received funding from the European Union’s Horizon 2020 research and innovation program under the Marie Sklodowska Curie grant agreement No 654609 (P.R-I). Q.Z. was funded by CSC and The University of Nottingham. W.W.C.T. was supported by the Croucher Foundation. M.A.S. is supported by Wellcome Investigator Award and core funding from Wellcome-CRUK to the Gurdon Institute. This work was supported by the Biotechnology and Biological Sciences Research Council [grant number BB/M001466/1] to R.A. and M.A.S.

## Author Contributions

P.R-I. designed and performed experiments including IF, scRNA-Seq, embryo dissections and wrote the paper; F.S. performed bioinformatics analysis; Q.Z. S.W., D.K. contributed to scRNA-Seq and IF. W.W.C.T. contributed with scRNA-Seq data; M.L. supervised scRNA-seq analysis; M.A.S. supervised scRNA-Seq experiments, advised on the project and wrote the paper. R.A. supervised the project, designed experiments, performed dissections and wrote the paper. All authors discussed the results and contributed to the manuscript.

